# Determining the impact of uncharacterized inversions in the human genome by droplet digital PCR

**DOI:** 10.1101/766915

**Authors:** Marta Puig, Jon Lerga-Jaso, Carla Giner-Delgado, Sarai Pacheco, David Izquierdo, Alejandra Delprat, Magdalena Gayà-Vidal, Jack F. Regan, George Karlin-Neumann, Mario Cáceres

## Abstract

Despite the interest in characterizing all genomic variation, the presence of large repeats at the breakpoints of many structural variants hinders their analysis. This is especially problematic in the case of inversions, since they are balanced changes without gain or loss of DNA. Here we tested novel linkage-based droplet digital PCR (ddPCR) assays on 20 inversions ranging from 3.1 to 742 kb and flanked by long inverted repeats (IRs) of up to 134 kb. Among these, we validated 13 inversions predicted by different genome-wide techniques. In addition, we have generated new experimental human population information across 95 African, European and East-Asian individuals for 16 of them, including four already known inversions for which there were no high-throughput methods to determine directly the orientation, like the well-characterized 17q21 inversion. Through comparison with previous data, independent replicates and both inversion breakpoints, we have demonstrated that the technique is highly accurate and reproducible. Most of the studied inversions are frequent and widespread across continents, showing a negative correlation with genetic length. Moreover, all except two show clear signs of being recurrent, and the new data allowed us to define more clearly the factors affecting recurrence levels and estimate the inversion rate across the genome. Finally, thanks to the generated genotypes, we have been able to check inversion functional effects in multiple tissues, validating gene expression differences reported before for two inversions and finding new candidate associations. Our work therefore provides a tool to screen these and other complex genomic variants quickly in a large number of samples for the first time, highlighting the importance of direct genotyping to assess their potential consequences and clinical implications.

## INTRODUCTION

During the last years a substantial amount of information has accumulated about different types of genomic changes, ranging from single nucleotide polymorphisms (SNPs) to more complex structural variants (SVs) (The 1000 Genomes Project Consortium 2015; Sudmant et al. 2015; Handsaker et al. 2015; Audano et al. 2019). However, inversions remain as one of the most difficult classes of variation to identify and characterize. Polymorphic inversions have been studied for many years and are known to have adaptive value and be associated with phenotypic effects in many species, including latitudinal clines in Drosophila, mimicry in butterflies, reproductive isolation in plants, adaptation to freshwater in stickleback fishes or different behaviors in sparrows, among many other examples (Umina et al. 2005; Joron et al. 2011; Lowry and Willis 2010; Jones et al. 2012; Thomas et al. 2008). In humans, hundreds of inversions have been predicted (Martínez-Fundichely et al. 2014; Puig et al. 2015a). However, for many of them it is not possible to detect reliably both orientations due to their balanced nature and the complexity of their breakpoints, and only a few have been analyzed in detail (Stefansson et al. 2005; Salm et al. 2012; González et al. 2014; Puig et al. 2015b; Giner-Delgado et al. 2019).

Approximately half of human inversions have IRs at their breakpoints (Martínez-Fundichely et al. 2014; Puig et al. 2015a), which can be up to hundreds of kilobases long. Therefore, short reads from next generation sequencing technologies (∼100-150 bp) (Sudmant et al. 2015; Hehir-Kwa et al. 2016; Collins et al. 2019), and even longer reads from single-molecule sequencing technologies (∼10 kb on average) (Huddleston et al. 2017; Shao et al. 2018; Audano et al. 2019) or paired-end mapping (PEM) data from large fragments (∼40-kb fosmid clones) (Kidd et al. 2008), are often not able to jump across the IRs at inversion breakpoints, precluding the detection of these inversions. New methods like single-cell sequencing of one of the two DNA strands by Strand-seq (Sanders et al. 2016; Chaisson et al. 2019) or generation of genome maps based on linearized DNA molecules labeled at particular sequences by optical mapping (Li et al. 2017; Levy-Sakin et al. 2019) have demonstrated their ability to detect inversions despite the presence of long IRs. Nevertheless, these techniques are not suitable for the analysis of large numbers of individuals.

The recent genotyping of 45 inversions in multiple individuals by a combination of inverse PCR (iPCR) and multiplex ligation-dependent probe amplification (MLPA) (Giner-Delgado et al. 2019) has increased extraordinarily the amount of data available on human inversions, although there are still limitations in the size of the IRs that can be spanned (up to ∼25-30 kb in this case). At the other end, low-throughput techniques like fluorescence in situ hybridization (FISH) have been used to study inversions with large IRs, but FISH is time consuming and can be applied only to inverted segments in the Mb scale where probes can be detected as separate signals (Antonacci et al. 2009; Salm et al. 2012). This leaves a set of potential inversions with IRs too large for iPCR-based techniques and inverted regions too small for FISH analysis that cannot be validated or genotyped in a large sample to determine their functional effects and evolutionary history.

Moreover, the great majority of inversions mediated by IRs have been shown to occur recurrently several times both within the human lineage and in non-human primates (Aguado et al. 2014; Vicente-Salvador et al. 2017; Giner-Delgado et al. 2019; Antonacci et al. 2009). Unlike inversions without IRs, which always have a single origin, these recurrent inversions tend to show low linkage disequilibrium (LD) to nearby nucleotide variants (Giner-Delgado et al. 2019), and as a consequence, they have been probably missed in existing genome-wide association studies (GWAS) of different phenotypes. In fact, several inversions have been associated to potential important effects on gene expression, phenotypic traits and disease susceptibility (de Jong et al. 2012; Myers et al. 2005; Zabetian et al. 2007; Webb et al. 2008; Salm et al. 2012; González et al. 2014; Puig et al. 2015b; Giner-Delgado et al. 2019). Inversion recurrence therefore highlights the need to expand the set of analyzed inversions by developing new tools able to genotype them directly through the order of the sequences around the breakpoints, rather than relying on linkage to other variants easier to genotype or computational methods based on SNP combinations (Ma and Amos 2012; Cáceres et al. 2012; Cáceres and González 2015).

In this context, droplet digital PCR (ddPCR) provides the opportunity to fill the void in inversion characterization by detecting linkage between amplicons located at both sides of the breakpoint repeats, making it possible to jump long genomic distances. The ddPCR^TM^ technology has already been used to detect copy number variation (Boettger et al. 2012), viral load (Strain et al. 2013), low-frequency transcripts (Hindson et al. 2013), rare mutations or cell free DNA (Olmedillas-López et al. 2017; Camunas-Soler et al. 2018) among other applications. Recently, it was shown that this technique could also be used to phase variants separated by up to 200 Kb (Regan et al. 2015), which has already been applied to fusion transcript detection (Hoff et al. 2016) or to phase deletions into haplotypes (Boettger et al. 2016). Here, we have developed new ddPCR assays to genotype quickly and reliably human polymorphic inversions flanked by large IRs, and thanks to the genotype data we demonstrate that most of these inversions are recurrent and that inversion alleles are associated to gene expression changes.

## RESULTS

### Inversion genotyping

To test the ddPCR application for inversion genotyping, we analyzed a representative sample of 20 inversions mediated by IRs of different sizes from the InvFEST database (Martínez-Fundichely et al. 2014). Due to the high error rate of inversion detection in repetitive sequences (Vicente-Salvador et al. 2017; Chaisson et al. 2019; Giner-Delgado et al. 2019), we selected well-supported inversions, excluding predictions on very complex regions full of segmental duplications (SDs) or with genome assembly gaps. In particular, 14 were initially predicted from fosmid PEM data in nine individuals (Kidd et al. 2008), although five had additional supporting evidence (Supplemental Table S1). The rest were validated inversions for which there was no available experimental genotyping method to study them in multiple individuals (the well-known 17q21 inversion or HsInv0573, HsInv0390, HsInv0290 and HsInv0786) (Stefansson et al. 2005; Beck et al. 2015; Feuk et al. 2005; Martin et al. 2004) and two control inversions already genotyped by iPCR-based assays (HsInv0241 and HsInv0389) (Aguado et al. 2014; Giner-Delgado et al. 2019) (Supplemental Table S1). Minimal inversion sizes range from 3.1 to 741.7 kb and IRs at breakpoints between 11.3 and 134 kb (Fig. 1A, Table 1), large enough to hinder inversion detection by conventional genome sequencing methods (see below).

**Figure 1.**
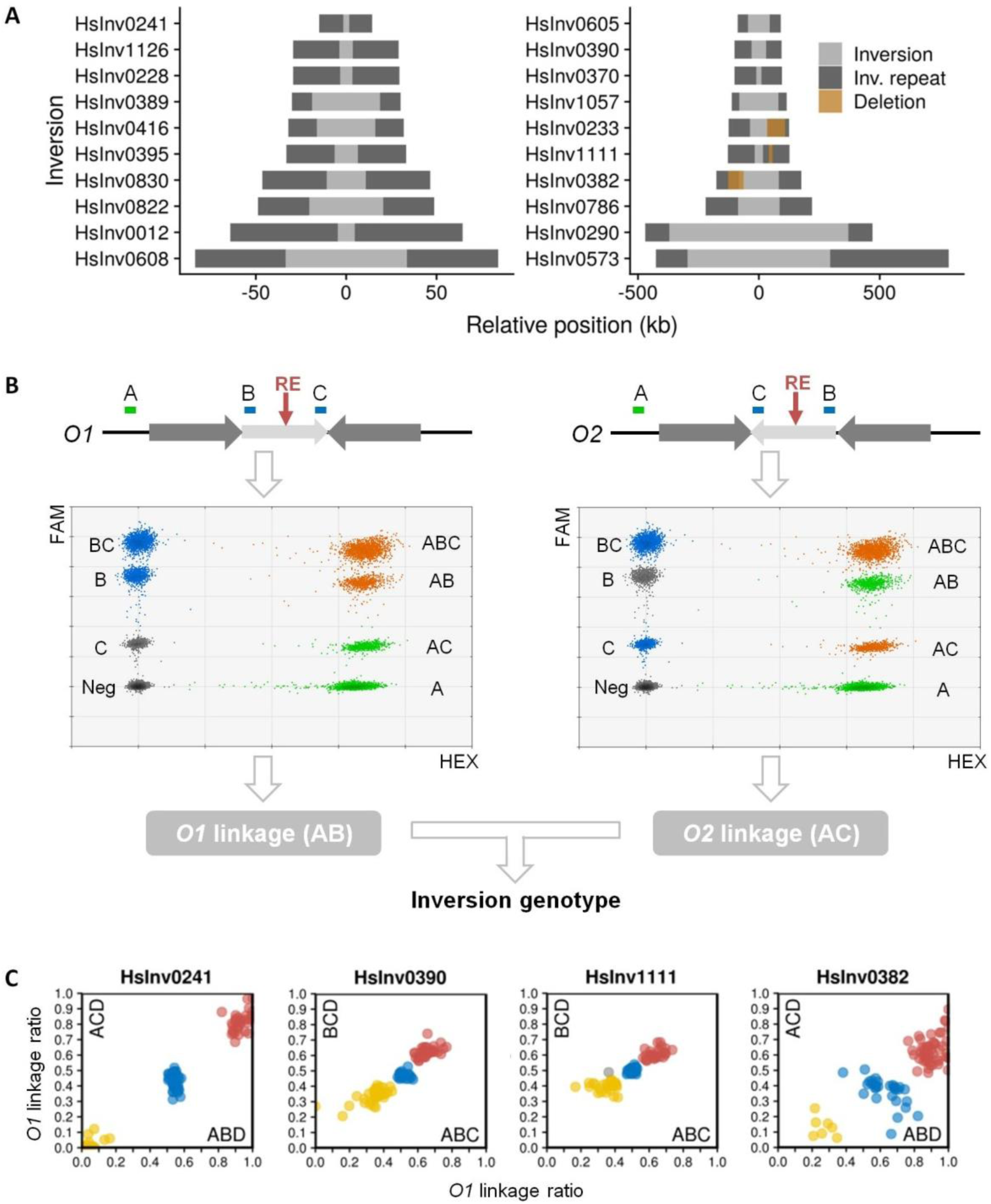
Inversion characterization by ddPCR genotyping. (*A*) Main features of the studied polymorphic inversions, with the inverted region shown in grey and the inverted repeats as dark boxes. Brown boxes denote deletions affecting inversion breakpoints. (*B*) Strategy used to genotype inversions. Dark grey arrows represent IRs at inversion breakpoints flanking the inverted region (light grey arrow), with ddPCR amplicons A, B and C shown on top. *O1* linkage (AB, left) and *O2* linkage (AC, right) were calculated separately from the triplex ddPCR results, ignoring respectively amplicons C or B. Colors of droplet clusters indicate if they are used to estimate the molecules containing only amplicon A (green), only the B or C internal amplicon (blue), two amplicons at the same time, either A and B or A and C (orange), or they are considered as negative (grey). (*C*) Plots of *O1* linkage ratios for both breakpoints of four inversions in 95 analyzed individuals. Colors indicate genotype groups (*O1/O1*, red; *O1/O2*, blue; *O2/O2*, yellow) and the grey dot is a sample with altered amplicon ratios in HsInv1111 ABC breakpoint. HsInv0241 clusters show the expected linkage ratios since DNA was digested to separate both breakpoints, whereas for the other inversions DNA digestion was not possible and homozygote and heterozygote clusters are closer. The effect of a large deletion within breakpoint AB in HsInv0382 can be seen as higher *O1* linkage ratio values in the horizontal axis.

**Table 1.**
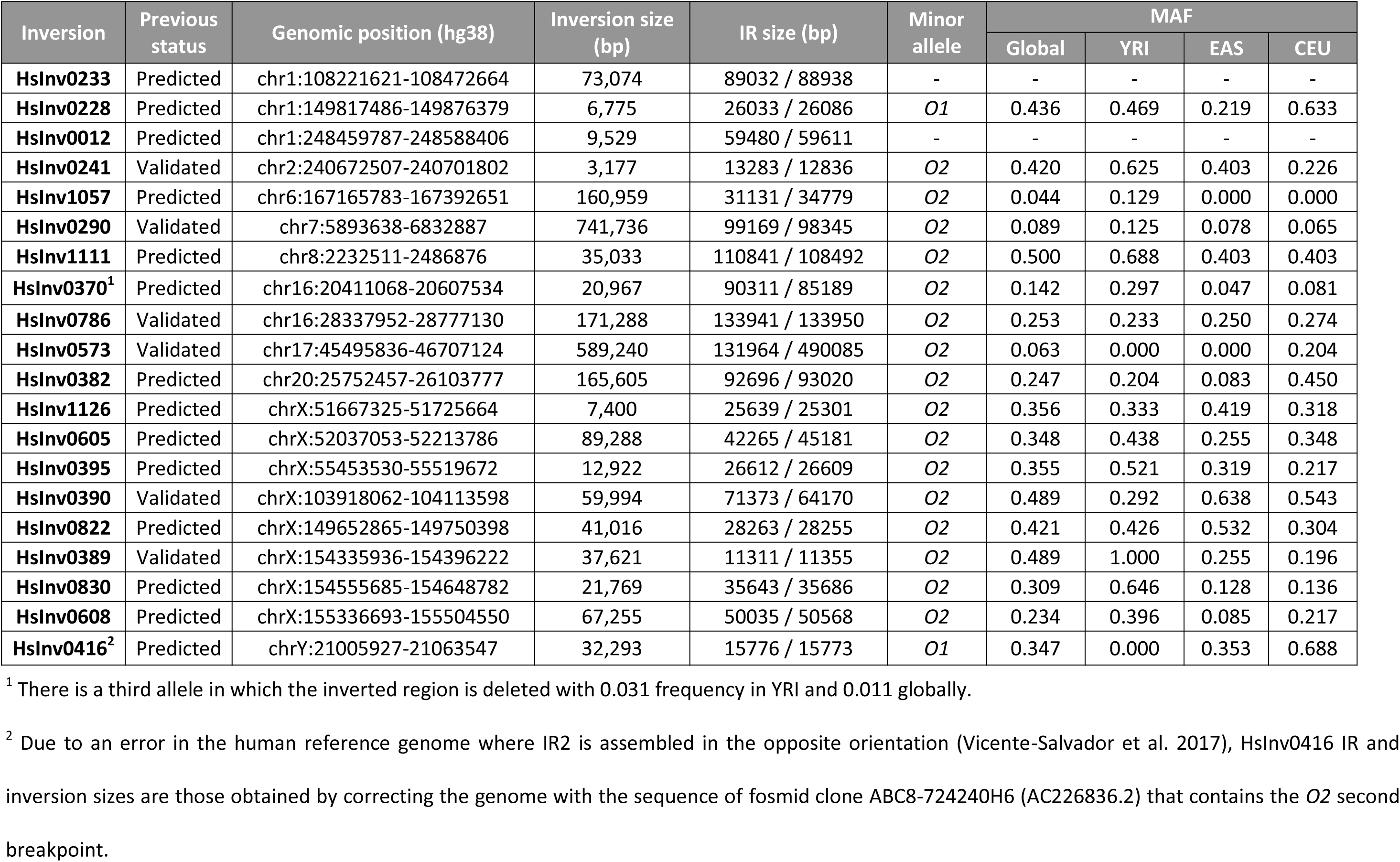
Inversion features and frequencies. Inversion sizes correspond to the distance between both inverted repeats (IR). The O1 allele corresponds to the orientation in the hg18 genome assembly, except for HsInv1111 in which it is the orientation in hg38 that represents the first complete sequence of the region. Inversion frequencies for the three analyzed populations together (global) and each population independently are those of the indicated global minor allele. AFR, Africans; EAS, East-Asians; EUR, Europeans.

| Inversion | Previous status | Genomic position (hg38) | Inversion size (bp) | IR size (bp) | Minor allele | MAF |  |  |  |
| --- | --- | --- | --- | --- | --- | --- | --- | --- | --- |
|  |  |  |  |  |  | Global | YRI | EAS | CEU |
| **HsInv0233** | Predicted | chr1:108221621-108472664 | 73,074 | 89032 / 88938 | - | - | - | - | - |
| **HsInv0228** | Predicted | chr1:149817486-149876379 | 6,775 | 26033 / 26086 | O1 | 0.436 | 0.469 | 0.219 | 0.633 |
| **HsInv0012** | Predicted | chr1:248459787-248588406 | 9,529 | 59480 / 59611 | - | - | - | - | - |
| **HsInv0241** | Validated | chr2:240672507-240701802 | 3,177 | 13283 / 12836 | O2 | 0.420 | 0.625 | 0.403 | 0.226 |
| **HsInv1057** | Predicted | chr6:167165783-167392651 | 160,959 | 31131 / 34779 | O2 | 0.044 | 0.129 | 0.000 | 0.000 |
| **HsInv0290** | Validated | chr7:5893638-6832887 | 741,736 | 99169 / 98345 | O2 | 0.089 | 0.125 | 0.078 | 0.065 |
| **HsInv1111** | Predicted | chr8:2232511-2486876 | 35,033 | 110841 / 108492 | O2 | 0.500 | 0.688 | 0.403 | 0.403 |
| **HsInv0370<sup>1</sup>** | Predicted | chr16:20411068-20607534 | 20,967 | 90311 / 85189 | O2 | 0.142 | 0.297 | 0.047 | 0.081 |
| **HsInv0786** | Validated | chr16:28337952-28777130 | 171,288 | 133941 / 133950 | O2 | 0.253 | 0.233 | 0.250 | 0.274 |
| **HsInv0573** | Validated | chr17:45495836-46707124 | 589,240 | 131964 / 490085 | O2 | 0.063 | 0.000 | 0.000 | 0.204 |
| **HsInv0382** | Predicted | chr20:25752457-26103777 | 165,605 | 92696 / 93020 | O2 | 0.247 | 0.204 | 0.083 | 0.450 |
| **HsInv1126** | Predicted | chrX:51667325-51725664 | 7,400 | 25639 / 25301 | O2 | 0.356 | 0.333 | 0.419 | 0.318 |
| **HsInv0605** | Predicted | chrX:52037053-52213786 | 89,288 | 42265 / 45181 | O2 | 0.348 | 0.438 | 0.255 | 0.348 |
| **HsInv0395** | Predicted | chrX:55453530-55519672 | 12,922 | 26612 / 26609 | O2 | 0.355 | 0.521 | 0.319 | 0.217 |
| **HsInv0390** | Validated | chrX:103918062-104113598 | 59,994 | 71373 / 64170 | O2 | 0.489 | 0.292 | 0.638 | 0.543 |
| **HsInv0822** | Predicted | chrX:149652865-149750398 | 41,016 | 28263 / 28255 | O2 | 0.421 | 0.426 | 0.532 | 0.304 |
| **HsInv0389** | Validated | chrX:154335936-154396222 | 37,621 | 11311 / 11355 | O2 | 0.489 | 1.000 | 0.255 | 0.196 |
| **HsInv0830** | Predicted | chrX:154555685-154648782 | 21,769 | 35643 / 35686 | O2 | 0.309 | 0.646 | 0.128 | 0.136 |
| **HsInv0608** | Predicted | chrX:155336693-155504550 | 67,255 | 50035 / 50568 | O2 | 0.234 | 0.396 | 0.085 | 0.217 |
| **HsInv0416<sup>2</sup>** | Predicted | chrY:21005927-21063547 | 32,293 | 15776 / 15773 | O1 | 0.347 | 0.000 | 0.353 | 0.688 |

<sup>1</sup> There is a third allele in which the inverted region is deleted with 0.031 frequency in YRI and 0.011 globally.
<sup>2</sup> Due to an error in the human reference genome where IR2 is assembled in the opposite orientation (Vicente-Salvador et al. 2017), HsInv0416 IR and inversion sizes are those obtained by correcting the genome with the sequence of fosmid clone ABC8-724240H6 (AC226836.2) that contains the O2 second breakpoint.

ddPCR technology allows us to quantify how close two independent sequences are within a DNA molecule based on their simultaneous amplification in a higher number of droplet partitions than expected by chance (Regan et al. 2015). Thus, for each inversion we designed three amplicons in the unique sequence outside the IRs (A or D) and at both ends of the inverted segment (B and C) (Fig. 1B). Then, we determined the percentage of DNA molecules supporting orientation 1 (*O1*) and orientation 2 (*O2*) from the linkage values between the amplicons outside and inside the inverted region, which correspond to the percentage of molecules containing both products (either A and B or C and D in the case of *O1* linkage, and A and C or B and D for *O2* linkage). Inversions were finally genotyped in each sample by the ratio between *O1* linkage and the total linkage for both orientations (*O1*+*O2* linkage), with ideal values for the three expected genotypes of 1 (*O1*/*O1*), 0.5 (*O1*/*O2*) and 0 (*O2*/*O2*). A different strategy was used for inversion 17q21, where all breakpoints contain variable repeat blocks >200 kb (Boettger et al. 2012; Steinberg et al. 2012), except for the AB breakpoint junction with a 132-kb repeat. Therefore, we measured only the *O1* linkage (AB) and compared it to a reference linkage (Ref) between two products located at the same relative distance in both orientations, resulting in *O1*/Ref linkage ratios of 1 (*O1*/*O1*), 0.5 (*O1*/*O2*) and 0 (*O2*/*O2*). In addition, to simplify the process and reduce costs, we developed triplex assays where the three amplicons from a given inversion are amplified simultaneously using different amounts of two probes labeled with FAM, which allows us to genotype an individual in a single reaction (Fig. 1B; Supplemental Table S2). These assays were tested and optimized in a small sample of 7-15 individuals, including those in which the inversions were predicted (see Methods).

The main problems in distinguishing ddPCR genotypes were related to inversion and IR size, variation in IR length and altered amplicon ratios. First, in small inversions linkage can be detected between A and C in *O1* chromosomes, or A and B in *O2* chromosomes, resulting in linkage ratios for homozygotes that depart from the 0/1 expected values. To solve this, whenever possible we separated both breakpoints by DNA digestion with a restriction enzyme that cuts the inverted sequence but not the IRs (Fig. 1B, Supplemental Table S2). In those inversions without a suitable restriction enzyme, linkage ratios for homozygotes were closer to those of heterozygotes, but we could still distinguish the three genotypes reliably (Fig. 1C). The exception was HsInv0012, in which the inverted region is so small compared to the IRs (Fig. 1A, Table 1) that amplicons B and C were equally linked to D in all samples and it was not analyzed further. Second, increasing distance between the ddPCR amplicons due to the IRs at the breakpoints reduces the linkage detected because there are less molecules long enough to contain both of them. In the inversions with the largest IRs, true linkage values become closer to those indicating no linkage, and the intrinsic variation of the technique can affect the final genotype. Thus, DNA length and quality limits the ability to resolve these inversion genotypes. Third, IR size variation caused by large polymorphic indels can change the distance between amplicons in different chromosomes of the population. These indels can also be moved to a different breakpoint by the inversion, altering more than one linkage value. Three of the studied inversions had polymorphic deletions within one of the IRs, but they only affected significantly the linkage in HsInv0382 (62-kb deletion within a 92.7-kb IR) and HsInv0233 (74.5-kb deletion within an 89-kb IR) (Supplemental Table S2). In HsInv0382 *O1/O2* individuals carrying the AB deletion, *O1* linkage is higher than *O2*, but three genotype groups can still be clearly separated (Fig. 1C). In HsInv0233, the large deletion in breakpoint CD combined with the relatively small inversion size leaves amplicons B and D at a similar distance in deleted *O1* chromosomes and in the inverted orientation, and it was not possible to assign genotypes with confidence (Supplemental Fig. S1). Finally, although all amplicons were carefully designed in unique single-copy sequences, we detected a few individuals that carry deletions or duplications that affect consistently one or more amplicons, which in the case of copy number increases makes it very difficult to interpret linkage values (Supplemental Table S3).

Next, we genotyped 19 inversions (excluding HsInv0012) in 95 individuals included in different genomic projects (The 1000 Genomes Project Consortium 2015; Lappalainen et al. 2013) with African (32 YRI), East Asian (EAS) (16 CHB and 16 JPT), and European (31 CEU) ancestry (Supplemental Table S4). All experiments were performed in duplicate, except for HsInv0389 that had been previously genotyped (Giner-Delgado et al. 2019), and five inversions for which the two breakpoints were analyzed independently (HsInv0241, HsInv0233, HsInv0382, HsInv0390 and HsInv1111). In HsInv0233, we identified different samples showing distinctive *O1/O1* and *O2/O2* genotypes with no signs of the large deletion and we consider this inversion validated, although we cannot genotype reliably all individuals. In general, genotypes for the rest of inversions were very clear and consistent between replicates. However, samples with problems (low linkage values in inversions with the largest IRs, intermediate linkage ratios that cannot be easily interpreted into genotypes, or altered amplicon ratios) were repeated several times. Final genotypes were called using a statistical clustering method that groups the *O1* linkage ratios of all the analyzed samples taking into account the information of every replicate (see Methods for details), which was especially important for inversions analyzed using undigested DNA (Fig. 1C). In total, we obtained 1635 genotypes for the 18 inversions with complete data (98.3%), and only 29 genotypes were not determined because of low linkage (13), inconclusive clustering results (12) or altered amplicon ratios due to CNVs (4) (Supplemental Table S3).

To assess the accuracy of the ddPCR results, we compared them with the published genotypes for all the samples for HsInv0241 and HsInv0389 (Giner-Delgado et al. 2019), plus a few from other inversions obtained by FISH or Southern blot (Supplemental Table S5). In addition, we genotyped by haplotype-fusion PCR (HF-PCR) (Turner et al. 2006) inversion HsInv0395 (90 CEU individuals) and HsInv0605 (8 individuals) with medium-sized IRs (26.6-45.2 kb) (Supplemental Fig. S2). In HF-PCR a fusion product is created from amplicons at both sides of a breakpoint, but it is difficult to set up and not very robust. Out of the 244 compared genotypes from 6 different inversions, there were only four discordant genotypes (1.6%) (Supplemental Table S5), which correspond to African individuals for HsInv0241 that appear to be *O2*/*O2* by iPCR, but are heterozygotes by ddPCR assays of both breakpoints. To check how common this discrepancy was, we genotyped the whole 30 YRI trios from the HapMap project (including 68 extra samples) (Supplemental Table S4) and only one child was discordant as well. Also, in three out of four candidates to have the same problem identified by SNP analysis from the LWK population, ddPCR genotypes were indeed *O1/O2* but *O2/O2* in iPCR (Supplemental Table S4). This suggests that there are some unknown variants affecting HsInv0241 *O1* chromosome detection by iPCR (Aguado et al. 2014; Giner-Delgado et al. 2019). Similarly, the 88 ddPCR 17q21 inversion genotypes agree with those inferred from the commonly-used tag SNPs (Steinberg et al. 2012). In contrast, in HsInv0786, 9 of the 81 genotypes (11.1%) computationally inferred from SNP genotype data (González et al. 2014) differ from ddPCR results (including eight heterozygotes misclassified as *O1*/*O1* and one *O1*/*O1* homozygote as heterozygote) (Supplemental Table S5), indicating errors in the imputation.

When ddPCR data were compared to recent inversion predictions in multiple human genomes with different techniques, quite contrasting results were obtained (Supplemental Table S1). Despite most of the inversions being relatively common, only HsInv0241, with some of the smallest IRs, was detected in one of the short-read analyses (Sudmant et al. 2015; Hehir-Kwa et al. 2016; Collins et al. 2019) and two more (HsInv0370 and HsInv1126) using long reads (Audano et al. 2019). On the other hand, two studies relying in a multiplatform strategy based mainly on Strand-seq (Chaisson et al. 2019) or Bionano optical maps (Levy-Sakin et al. 2019) identified 14 inversions each, with 9 detected by both (Supplemental Table S1). Although in most cases genotypes were not provided, based on the presence or not of the inverted allele of the identified inversions, short reads (Sudmant et al. 2015) and long reads (Audano et al. 2019) showed the lowest performance (with respectively 66% and 83% error rate). The Bionano analysis missed the inverted alleles in 23% of individuals, and the multiplatform strategy genotyped correctly 10 of 11 comparable inversions in the only individual in common (9% error rate). This emphasizes the amount of inversion information still missing from the studied genomes and the need of specific methods for the analysis of IR-mediated inversions.

With regard to the distribution and frequency of these inversions (Table 1), all are in Hardy-Weinberg equilibrium in the analyzed populations and most of them are widespread in the three continents with global minor allele frequencies (MAF) >0.1. The only exceptions are HsInv1057, found exclusively in Africans, the 17q21 inversion, detected only in Europeans where the frequency of the H2 allele is known to be higher (Stefansson et al. 2005; Steinberg et al. 2012; Alves et al. 2015), and HsInv0290 with low frequency in the three populations (0.065-0.125). Several inversions also have variable frequencies between continents, with chr. X inversions HsInv0830 and HsInv0389, and HsInv0416 in chr. Y showing the highest F_ST_ values, although only those of HsInv0389 are within the top 5% of the expected distribution (Supplemental Table S6) (Giner-Delgado et al. 2019). As observed before (Giner-Delgado et al. 2019), we found a negative correlation between inversion size and MAFs in each population either for the inversions analyzed here only or all inversions together (Fig. 2). These correlations tend to be higher with the inversion genetic length than the physical length (Fig. 2), which suggests that in large inversions negative selection against the generation of unbalanced gametes by recombination in heterozygotes (Hoffmann and Rieseberg 2008; Kirkpatrick 2010) might be an important factor in determining their frequency.

**Figure. 2.**
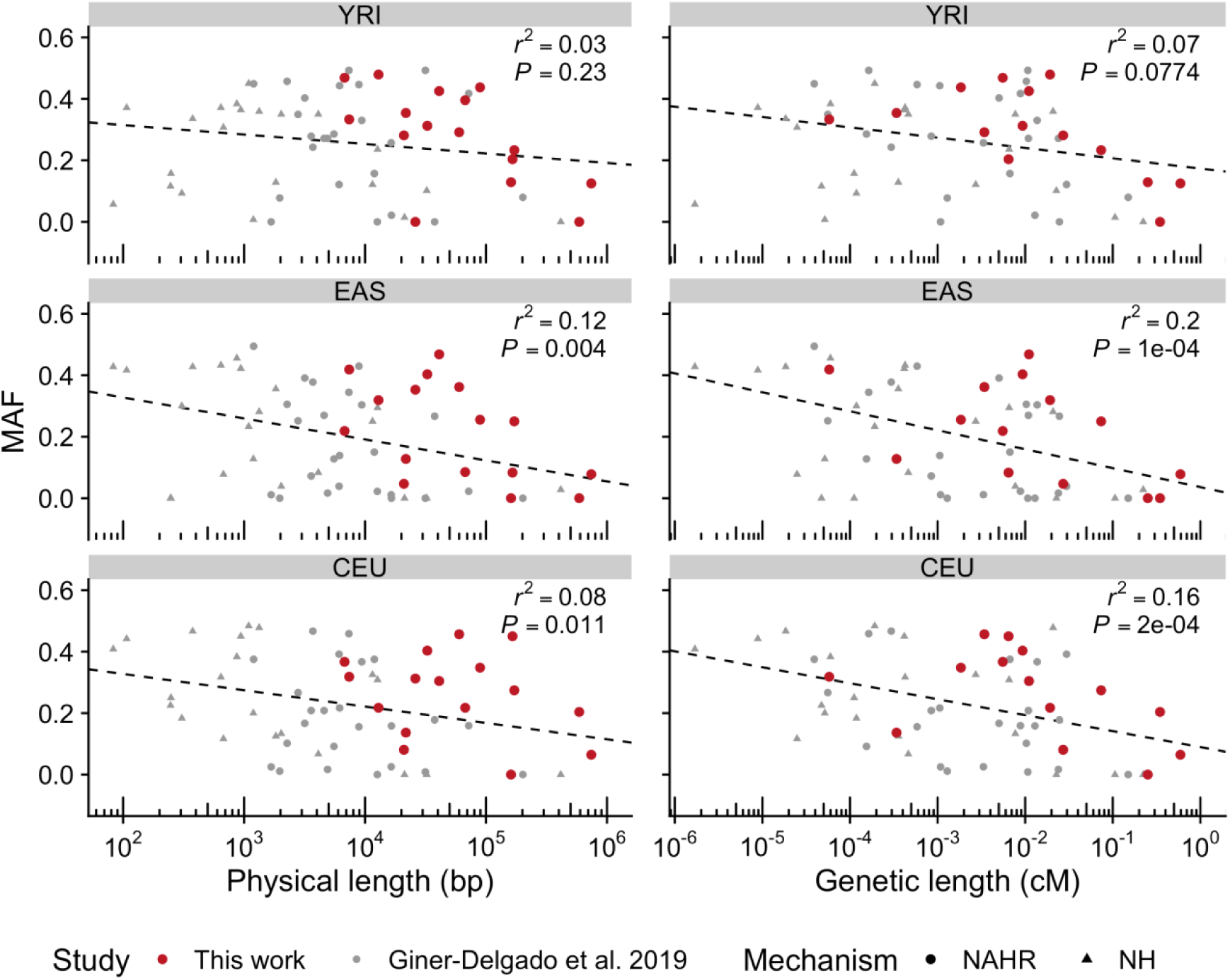
Correlation between inversion size and frequency. Negative correlations between the logarithm of the minimal physical or genetic length and minor allele frequency (MAF) of the inversion in three populations with African (YRI), East-Asian (EAS) and European (CEU) ancestry are observed for the 16 ddPCR-analyzed inversions (in red) and 45 polymorphic inversions described in Giner-Delgado et al. (2019) (in grey), including inversions generated by non-homologous mechanisms (NH) or non-allelic homologous recombination (NAHR).

### Nucleotide variation analysis

We used the accurate genotypes of 15 inversions to check the linkage disequilibrium (LD) with 1000 Genome Project (1000GP) nucleotide variants (92 individuals in common), excluding HsInv0416 in chr. Y without 1000GP data and HsInv0241 and HsInv0389 already analyzed in a larger number of individuals (Giner-Delgado et al. 2019). Only population-specific inversions 17q21 and HsInv1057 have tag SNPs in complete LD (*r^2^* = 1), while for the rest of inversions the maximum LD (*r^2^*) values range between 0.21 and 0.79 and just five of them have tag SNPs (*r^2^*> 0.8) in at least one of the populations (CEU, EAS or YRI) (Fig. 3A, Supplemental Table S7). We also classified the SNPs within the inverted region as shared between orientations, private to one of them or fixed (i.e. in complete LD) (Fig. 3A, Supplemental Table S7). The 17q21 and HsInv1057 inversions show no reliable shared variants, consistent with the inhibition of recombination across the entire inversion length. In contrast, shared variants represent an important fraction (6-48%) of the variation within the entire length of the remaining inversions (Supplemental Fig. S3), a pattern suggestive of several inversion events on different haplotypes.

**Figure 3.**
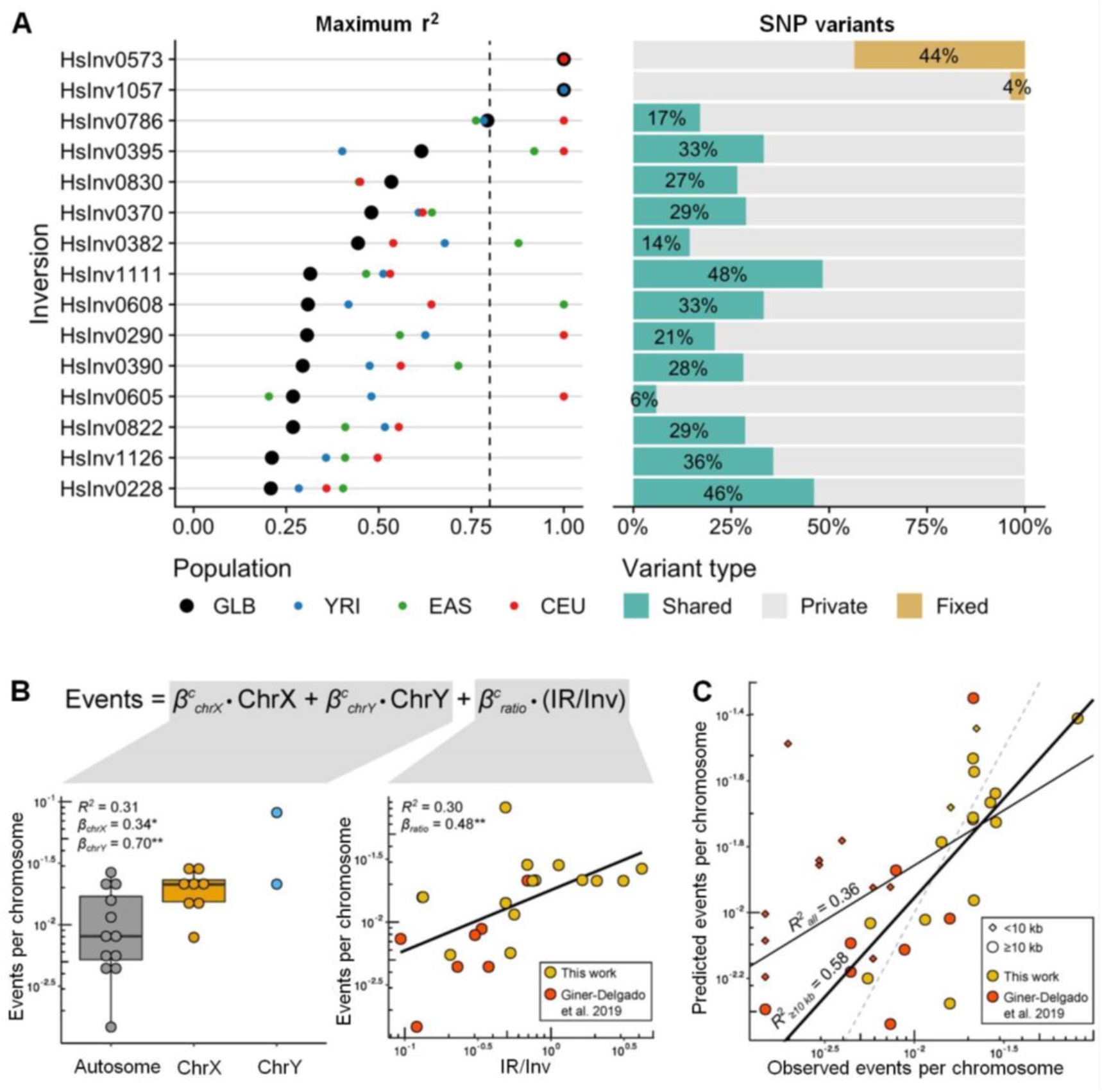
Nucleotide variation analysis of newly genotyped autosomal and chr. X inversions. (*A*) Left, maximum LD (*r^2^*) between 1000GP variants located 1 Mb at each side and inversion alleles in all individuals together (black dots) and different populations (colored dots). Right, proportions of SNPs classified as fixed (yellow), shared between orientations (green) or private of one orientation (grey) within each inversion. (*B*) Correlation between the logarithm-transformed estimated number of independent inversion events per chromosome and chromosome type (left) and the IR/inverted region (Inv) size ratio (right) calculated with 22 inversions >10 kb using data from this work (yellow dots) and Giner-Delgado et al. (2019) (orange dots). (*C*) Adjustment of the observed recurrence events with the expected number calculated by applying the developed model. Number of events is underestimated in small inversions (<10 kb; diamonds), which results in a lower *R*^2^ value for all inversions (all, thin black line) than for those greater than 10 kb (≥10 kb, thick black line). Dashed line represents the 1:1 equivalence.

Next, we estimated the independent inversion events by phasing inversion genotypes into 1000GP haplotypes of the inverted region (Fig. 4). While in the two inversions with perfect tag SNPs all *O2* haplotypes cluster together, in the other inversions several different clusters of similar haplotypes containing both orientations can be identified, which are typical of recurrent inversion and re-inversion events (Giner-Delgado et al. 2019). We identified between 2 and 5 inversion events in the 14 new recurrent inversions analyzed here and a total of 33 additional inversion events, with 27 shared by more than one individual (Supplemental Table S8). This suggests that they are real evolutionary events rather than phasing errors or new variants generated in the lymphoblastoid cell lines (LCLs) used as DNA source. However, recurrence quantification might be complicated by inversion and SNP phasing errors and recombination between haplotypes. The exception is HsInv0416, in which the *O1* and *O2* distribution in the known phylogeny of chr. Y haplogroups (Poznik et al. 2016) clearly supports four independent inversion events (Supplemental Table S8) and results in an inversion mutation rate of 1.29 x 10^-4^ inversions per generation, very similar to that previously described for another Chr. Y inversion (0.53 x 10^-4^ inversions per generation) (Giner-Delgado et al. 2019).

**Figure 4.**
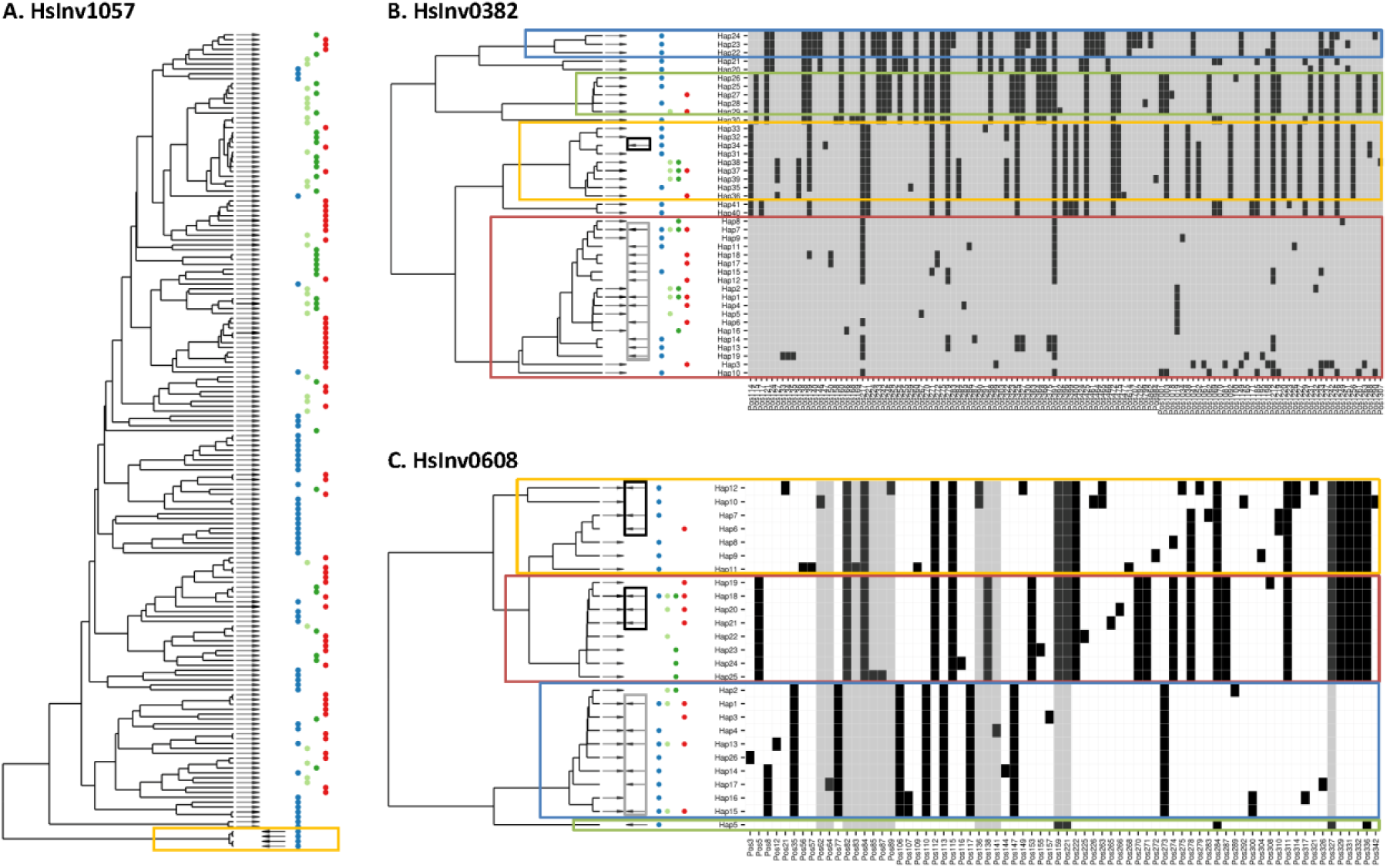
Estimation of the number of inversion events from inverted region haplotypes. Each inversion was analyzed using integrated haplotype plots (iHPlots) (Giner-Delgado et al. 2019), with the tree indicating the relationship between the different haplotypes, the rightwards (*O1*) and leftwards (*O2*) arrows the orientations observed for each haplotype, and dots the populations where each haplotype has been found (blue for YRI, light and dark green for CHB and JPT, and red for CEU). Inverted region haplotypes are represented by the variable positions (see Methods for variant selection) with different colors indicating the two alleles (white, ancestral; grey, hg19 reference; black, derived/alternative), and colored boxes showing the main differentiated haplotype clusters. For the unique inversion (*A*) only the first part of the plot is shown with a yellow box highlighting the single group of tightly-clustered inverted haplotypes. In the recurrent inversions (*B*, *C*), the orientation of the haplotypes of what we considered the original inversion event are included in a grey box and any additional inversion event in a black box (one in HsInv0382 and two in HsInv0608).

We also investigated the effect of different parameters on the observed recurrence levels by combining all the inversions mediated by NAHR analyzed here and in Giner-Delgado et al. (2019). Focusing only on the 22 inversions >10 kb, which have more resolution to detect recurrence events, the only significant variables were the ratio between IR and inversion sizes (IR/Inv ratio) and chromosome type (autosome, chr. X or chr. Y), with IR/Inv ratio and sex chromosomes being positively correlated with the number of inversion events (Fig. 3B). The resulting model fits very well the real data (*R^2^* = 0.58), although there is a clear underestimation of recurrence events in smaller inversions (<10 kb) because the lower number of SNPs reduces the ability to distinguish haplotype clusters (Fig. 3C).

### Functional effects of inversions

The functional consequences of the genotyped inversions were first investigated through their association with available LCL gene expression data (Lappalainen et al. 2013) of 59 CEU and YRI samples in common. We identified two inversions associated to gene-expression changes in this small sample (Fig. 5A, Supplemental Table S9). In particular, inversion 17q21 is the lead variant (*r^2^*≥ 0.8 with top eQTLs) for an antisense RNA and three pseudogenes at the gene level, as well as for specific transcripts from protein-coding genes *KANSL1* and *LRRC37A2* plus other pseudogenes and non-coding RNAs (Supplemental Table S9), which indicates a pervasive effect of the two inversion haplotypes on gene expression. To increase the power to detect associations, we extended the analysis to a larger sample of 387 Europeans for five inversions whose genotypes could be inferred through perfect tag SNPs (*r^2^* = 1) in the CEU population (Fig. 3A). We found that all of them were significantly associated to expression changes in 42 genes and 103 transcripts (Fig. 5A, Supplemental Table S9), including multiple additional genes for inversion 17q21. In addition, HsInv0786 appeared as potential lead eQTL for gene *NFATC2IP*, and *APOBR*, *IL27* and *EIF3C* transcripts.

**Figure 5.**
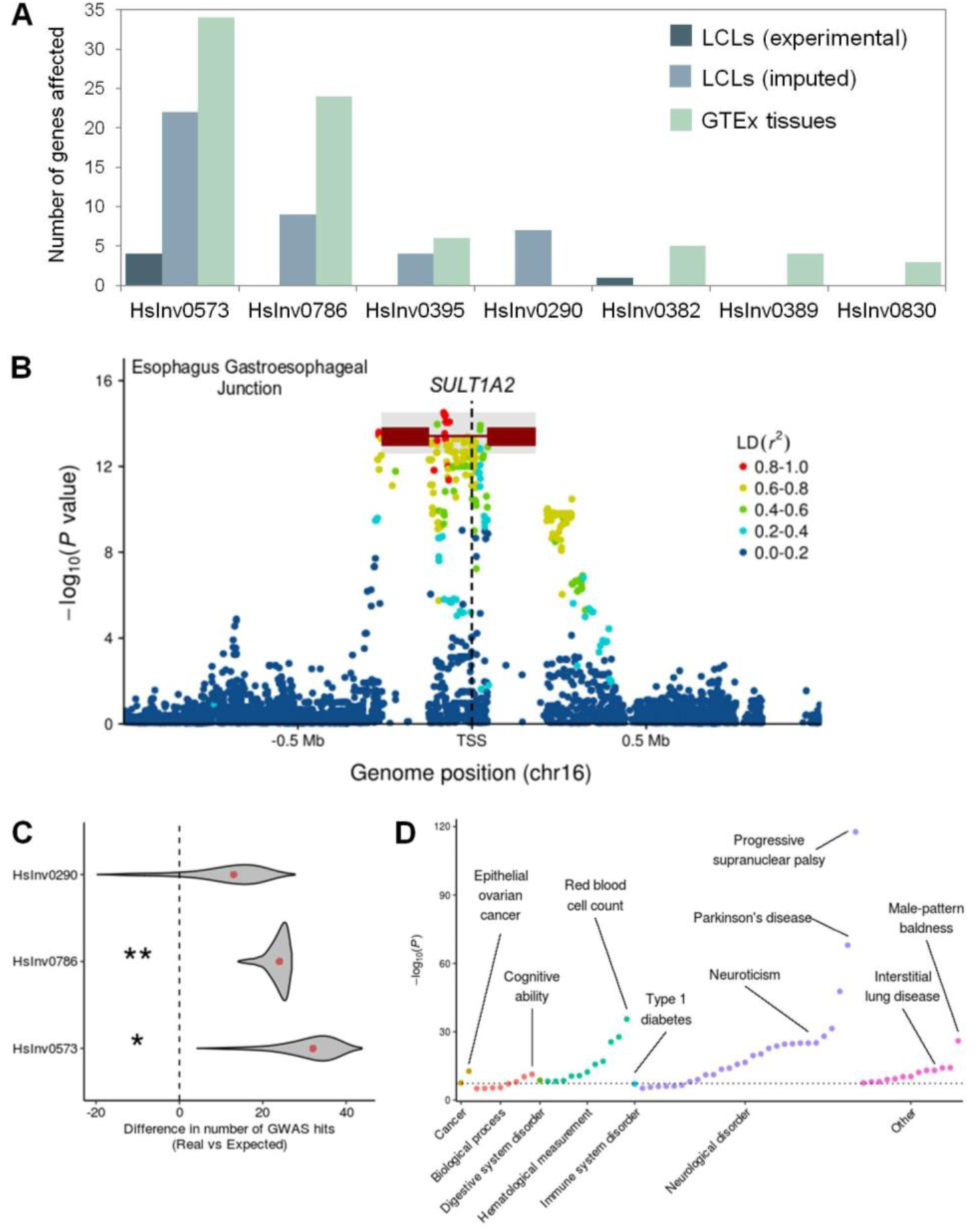
Inversion effects on gene expression and phenotypic traits. (*A*) Number of significant associations at the gene level in each of the differential expression analyses. (*B*) Manhattan plot for *cis*-eQTL associations reported in GTEx project of gene *SULT1A2*, showing inversion HsInv0786 (dark red line with rectangles representing the IRs) as potential lead variant. HsInv0786 eQTL *P* values and LD with neighboring variants were calculated by permuting samples with the same ancestry proportions as in GTEx samples (see Methods) and the *P* value imputation range is shown in grey. (*C*) Enrichment of GWAS Catalog signals within individual inversions measured by the deviation in observed minus expected value. Density distribution represents the 95% one-sided confidence interval with the median indicated by a red dot (one-tailed permutation test: **, *P* < 0.01; *, *P* < 0.05). (D) Phenome-wide association study for inversion 17p21. Significant reported traits (*P* < 10^-5^) were grouped by categories according to terms defined by GWAS Catalog ontologies. Dotted line indicates genome-wide significance threshold (*P* = 5 × 10^-8^).

Since more than 60% of genes inside or around our inversions (±1 Mb) are not expressed in LCLs, we explored their effects in other tissues using eQTL information from the GTEx Project (GTEx Consortium 2017). We imputed inversion association *P* values by estimating LD patterns between inversion alleles and SNPs identified as eQTLs in GTEx (see Methods). We found 73 genes linked to 6 inversions in different tissues, although the majority of associations involve inversions 17q21 and HsInv0786, whose effects are easier to infer due to their high LD with neighboring eQTLs (including 37 genes for which the inversions were potential lead variants) (Fig. 5A-B, Supplemental Table S10, Supplemental Fig. S4). However, the low LD patterns with SNPs due to high recurrence levels of most analyzed inversions prevents inferring their contribution to gene expression variation.

We also checked whether inversions account for phenotypic variation by using the GWAS Catalog information (MacArthur et al. 2017). We found a 2.8-fold significant enrichment (*P* = 0.026) of GWAS Catalog signals within inversion regions compared to other genomic regions of similar size, with 17p21 (*P* = 0.028), HsInv0786 (*P* = 0.009) and HsInv0290 (*P* = 0.157) apparently driving this enrichment (Fig. 5C). When we mapped GWAS phenotypes to genes located within 150 kb of the analyzed inversions, several of them showed an enrichment of GWAS-reported genes for certain traits (Supplemental Table S11), such as brain-related traits (Parkinson’s disease, neuroticism, intelligence and cognition) for 17p21 and immunological disorders (Crohn’s disease, ulcerative colitis and type 1 diabetes) and body mass characteristics for HsInv0786, indicating their potential roles on these diseases. For seven inversions in high LD (*r*² ≥ 0.8) with SNPs in at least one population, we also looked for GWAS hits associated to those SNPs in the corresponding population or continent (Supplemental Table S12). The 17p21 inversion has already been linked to many traits (Puig et al. 2015a) and a total of 64 potential associations with *P* < 10^-6^ from 35 studies were found in this analysis (Fig. 5D), including lung function, neurological traits or disorders, ovarian cancer, blood profiles and diabetes, which illustrates the pleiotropic consequences of this inversion. HsInv0786, which was associated to an asthma and obesity phenotype (González et al. 2014), can also be associated to several pediatric autoinmune diseases (Li et al. 2015) and the presence of type 1 diabetes autoantibodies (Plagnol et al. 2011) among other traits, suggesting a role in the immune system that may be related to the observed expression changes in genes like *IL27* and *NFATC2IP* (Supplemental Tables S9-S10).

## DISCUSSION

The study of polymorphic inversions, and especially those flanked by large IRs, has lagged behind because of the lack of high-throughput techniques able to detect them. Here, we have taken advantage of ddPCR ability to measure linkage between sequences at relatively long distances to genotype inversions with IRs up to ∼150 kb in a single reaction in an efficient, reliable and reproducible way. Using this method we have validated 13 inversion predictions and generated new experimentally-resolved genotypes for 16 inversions, most of which are analyzed here in detail for the first time. In comparison with the set of 24 inversions flanked by IRs recently analyzed (Giner-Delgado et al. 2019), on average our 16 inversions are longer (138.9 kb versus 20.7 kb) and have much bigger IRs (74.2 kb versus 6.9 kb), filling a void in the study of human variation. Almost all these inversions are missed by next-generation sequencing with short or long reads (Supplemental Table S1). In addition, although recent studies using other methods like Strand-seq or Bionano (Chaisson et al. 2019; Levy-Sakin et al. 2019) have been able to identify 95% of them (Supplemental Table S1), an acceptable genotyping accuracy (>90%) is only obtained by combining different techniques, which complicates the analysis of a large number of individuals. Therefore, the novel ddPCR application provides a very valuable and powerful resource for the targeted characterization of inversions and complex SVs, including accurate information on other associated variants in the region (like indels or CNVs) (Supplemental Table S3).

In fact, half of inversions mediated by inverted SDs in InvFEST (Martínez-Fundichely et al. 2014) have IRs between 25 and 150 kb long that could be analyzed using the ddPCR technology. The main limitations are restricted to extremely long IRs (>100 kb), small inversions (<5-10 kb) with relatively long IRs where breakpoints cannot be separated by restriction-enzyme digestion, and CNVs altering the distance between amplicons. We have overcome these problems by using good-quality high-molecular-weight DNA and a clustering tool that allows us to distinguish the three genotype groups independently of the magnitude of the linkage ratio differences. In the near future, the possibility to work with even longer DNA molecules and to interrogate several inversions simultaneously by multiplexing amplicons labeled with different fluorochromes will expand the range of studied inversions and reduce the costs, making easier to undertake more ambitious ddPCR genotyping projects in higher numbers of samples.

Consistent with previous results (Giner-Delgado et al. 2019), we have shown that the vast majority of inversions mediated by IRs are generated multiple times on different haplotypes by NAHR and cannot be easily imputed from SNP data. For example, in HsInv0786, half of individuals incorrectly genotyped using SNP information (11.1%) (González et al. 2014), carry a recurrent chromosome with unexpected orientation according to its haplotype. Also, inversion recurrence or some other mechanism able to exchange variants between the two IRs had been already suggested for HsInv0390 (Beck et al. 2015), HsInv0830 (Aradhya et al. 2001) and HsInv0290 (Hayward et al. 2007) regions. The only exceptions are the two inversions with more restricted geographical distributions: HsInv1057 (found exclusively in Africans) and 17q21 (with a higher frequency in Europeans compared to all other populations) (Stefansson et al. 2005; Steinberg et al. 2012; Alves et al. 2015). This indicates that although the 17q21 and a few other unique inversions can be indirectly imputed by tag SNPs, the only way to genotype accurately recurrent inversions is to interrogate experimentally the sequences at both sides of the breakpoint with techniques like the one developed here.

Actually, we have found similar levels of recurrence for the previous smaller inversions (Giner-Delgado et al. 2019) and the longer inversions studied here, showing that it is a general characteristic of inversions flanked by IRs. Moreover, the higher resolution provided by the longer inversions has allowed us to estimate more precisely the number of recurrence events and determine the main factors affecting recurrence. Specifically, the chromosome type and IR/inversion size ratio together explain a very significant part (up to 58%) of the variance in recurrence levels between inversions. This fits well the expectations, since repeat length and distance has been found to affect the generation of recurrent pathological rearrangements (Liu et al. 2011), suggesting that the closer and longer the IRs are, the more likely they are to pair and recombine. In addition, the inversion causing hemophilia A is known to occur much more frequently in the male germ line, probably due to the increased NAHR within the single chr. X copy (Antonarakis et al. 1995). According to the estimated values for two chr. Y inversions, the model results in predicted NAHR mutation rates for the analyzed autosomal inversions of 0.9-4.4 × 10^-5^ inversions/generation and of 1.9-7.4 × 10^-5^ inversions/generation for chr. X inversions, which illustrate the importance of this phenomenon and its potential impact in the genome. However, other factors, like DNA 3D conformation or recombination motifs, might also affect recurrence.

On the other hand, the newly-generated genotypes have allowed us to carry out the first complete analysis of the potential functional consequences of the majority of these inversions. By taking advantage of different available datasets and analysis strategies, we were able to associate eight inversions with gene-expression changes across different tissues. In particular, the inversions with the largest effects were 17q21 and HsInv0786, which were the top eQTLs for many genes. In this case, our analyses confirmed most expression changes already reported for 17q21 in blood, cerebellum and cortex (de Jong et al. 2012) or for HsInv0786 in blood and LCLs (González et al. 2014) and estimated more precisely the real contribution of the inversions. Moreover, we identified expression differences in additional genes and tissues never examined before (Supplemental Table S10, Supplemental Fig. S4. We also found that the two same inversions are also enriched in GWAS signals, including certain phenotypic categories such as neurological traits or immunological disorders, and they are in LD with multiple variants associated to different phenotypes. Thus, these results show that inversions could have important effects on both gene expression and clinically relevant disorders and uncover other interesting candidates for further study.

However, the inversion functional analysis has two main limitations. First, the small number of genotyped individuals with expression data (59) results in low statistical power and only large differences can be detected, especially for low-frequency inversions. This is exemplified by the additional expression associations identified for 17q21 and four other inversions when genotypes were imputed in additional European individuals (387). Second, for many inversions the lack of LD with neighboring variants that can be used as a proxy does not allow us to infer reliably their association with gene expression in other tissues or with phenotypic traits from non-genotyped individuals. Therefore, we are probably missing a significant fraction of inversion effects, especially those of modest sizes, emphasizing the need to genotype these inversions in a larger set of individuals.

Another important effect of inversions is the generation of aberrant chromosomes by recombination crossovers within the inverted region in heterozygotes (Hoffmann and Rieseberg 2008; Kirkpatrick 2010). By extending the analysis to a different set of inversions, here we have reinforced the idea that there is a negative correlation between the frequency and genetic length of inversions related to their negative impact in fertility (Giner-Delgado et al. 2019). In that sense, it is noteworthy that some of the longer inversions, such as 17q21 (589 kb) and Hsinv0786 (171 kb), are the best examples of inversions with functional consequences at different levels. This suggests that some of their effects on gene expression could compensate the potential negative costs associated to the longer inversions, and it has already been proposed that the 17p21 inversion has been positively selected in European populations through increased fertility in carrier females (Stefansson et al. 2005).

Finally, polymorphic inversions could also predispose to other pathological SVs in the region, due either to recombination problems in heterozygotes or changes in the orientation of repeats (Puig et al. 2015a). Recently, a complete catalogue of inversions in nine individuals has shown that a high proportion of them overlap the critical regions of microdeletion and microduplication syndromes (Chaisson et al. 2019). However, these inversions tend to be mediated by large and complex repeats and are difficult to characterize with simple methods. In our case, 7 of the 8 chr. X inversions analyzed are located in regions where additional SVs disrupting genes and resulting in disease have been described (Supplemental Table S13). Some of these diseases involve recurrent mutations mediated by repeats within the polymorphic inversions, like the inversion causing Hemophilia A and HsInv0608 (Antonarakis et al. 1995) or the deletion causing incontinentia pigmenti and HsInv0830 (Aradhya et al. 2001). In HsInv0389 and HsInv0390, DNA polymerase stalling during replication of the repeats has been associated to different duplication-inverted triplication-duplication (DUP-TRP/INV-DUP) rearrangements that affect dose-sensitive genes (*MECP2* and *PLP1*, respectively) (Carvalho et al. 2011; Beck et al. 2015). Similarly, different deletions with one breakpoint mapping nearby or within one of the HsInv1126 IRs and affecting the inversion region and the ∼310 kb separating it from HsInv0605 have been associated to X-linked intellectual disability (Grau et al. 2017). Interestingly, within the 6-Mb chr. Xq28 region near the telomere, a total of six genomic disorders caused by the previous and several other rearrangements overlapping or in close proximity to polymorphic inversions HsInv0822, HsInv0389, HsInv0830 and HsInv0608 have been described (Bondeson et al. 1995; Small et al. 1997; Clapham et al. 2012; Aradhya et al. 2001; Fusco et al. 2012; Li et al. 2015), making this region a possible hotspot for genome reorganization (Fig. 6). The new ddPCR application thus offers now the opportunity to study easily these inversions in parents of patients and determine their role in the generation of pathological variants, contributing to having a more complete picture of the impact of inversions in the human genome.

**Figure 6.**
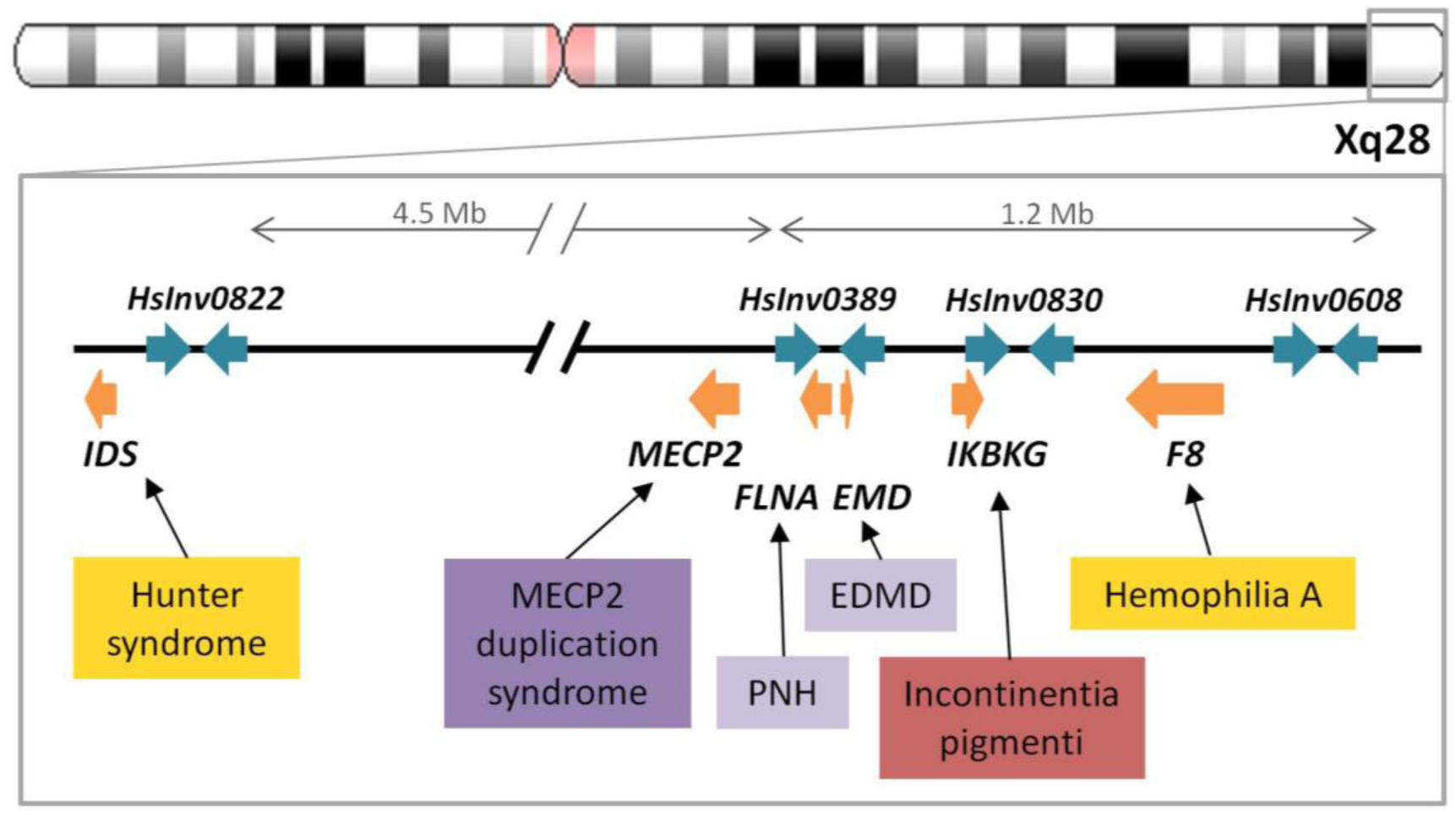
Polymorphic inversions in chromosome region Xq28 and diseases caused by structural rearrangements. Polymorphic inversion IRs are represented as blue arrows and genes as orange arrows. Colored boxes indicate diseases associated to different types of recurrent pathological rearrangements: inversions (yellow), deletion (red), and complex events resulting in deletion (light purple) or in a duplication-inverted triplication-duplication structure (DUP-TRP/INV-DUP) (dark purple). EDMD, Emery-Dreifuss muscular dystrophy; PNH, periventricular nodular heterotopia.

## METHODS

### Human samples and DNA isolation

Genomic DNA of 95 unrelated human samples (Supplemental Table S4) was isolated from ∼20-ml culture of an Epstein-Barr virus-transformed B-lymphoblastoid cell line of each individual (Coriell Cell Repository, Camden, NJ, USA) grown according to the recommended procedures. To obtain high-molecular-weight DNA, the cell pellet was resuspended in extraction buffer (10 mM Tris-HCl pH 8, 10 mM EDTA pH 8, 150 nM NaCl, 0.5% SDS) and incubated overnight with slow rotation at 37 °C with RNase cocktail (Invitrogen) and 100 μg/ml Proteinase K (Invitrogen). Four purification steps with one volume of TE-equilibrated phenol pH 7.9 (twice), phenol:chloroform:isoamyl alcohol pH 7.9, and chloroform:isoamyl alcohol were performed by mixing by rotation until an emulsion was formed, and then centrifuging at 5,000 x g for 10-15 min to separate organic and aqueous phases. All steps that involved handling of the DNA sample were done pipetting gently with wide-bore tips. Finally, DNA was precipitated by adding 0.1 volumes of 3 M sodium acetate and 2 volumes of absolute ethanol, centrifuged, washed with 70% ethanol, and resuspended in 100-300 μl of water. DNA was stored at 4 °C, which we observed that preserved DNA integrity over time better than freezing. Identity of all the isolated DNAs was confirmed by microsatellite analysis.

### Droplet digital PCR (ddPCR)

Inversion genotyping by ddPCR assays was carried out by quantitative amplification of three different products simultaneously with six different primers and three fluorescent probes (Supplemental Table S14) in aqueous droplets within an oil phase (emulsion PCR). Since the QX200^TM^ ddPCR system can detect only two colors, we labeled the probe in the amplicon common to both inversion alleles (A in an ABC experiment) with HEX and the other two (B and C) with FAM and used a lower concentration of one of the FAM probes to separate the clusters of droplets including the different amplicon combinations (Fig. 1B). All primers and probes were tested in duplex experiments before optimizing the triplex reactions. Final ddPCR reactions were prepared in a total volume of 22 μl in a 96-well plate with 450 nM-2.5 μM of each primer, 75-275 nM of each probe, 1x ddPCR Supermix for Probes (No dUTP) (Bio-Rad) and 50 ng of genomic DNA. DNA samples were always handled with wide-bore tips and the ddPCR mix was mixed by gently pipetting up and down 5-10 times to avoid breaking the DNA molecules. Droplet generation, thermal cycling and fluorescence reading were performed as explained before (McDermott et al. 2013) using the Bio-Rad QX200^TM^ Droplet Generator, C1000^TM^ or T100^TM^ thermal cyclers and QX200^TM^ Droplet Reader. QuantaSoft^TM^ version 1.7.4 software (Bio-Rad) was used to analyze each sample twice, every time ignoring one of the FAM products and counting droplets with that product the same as negatives (Fig. 1B). The linkage between the two targeted amplicons (percentage of DNA molecules containing both amplicons) was obtained as the excess of double-positive droplets over what is statistically expected (Regan et al. 2015). From these values we could calculate the total linkage and the *O1* and *O2* linkage ratios that allow us to genotype inversions. For inversions with restriction enzyme targets inside the inverted region but not within the IRs at the breakpoints, DNA was digested prior to ddPCR quantification. This helps to: (1) Separate both breakpoints and avoid the detection of linkage between undesired products due to their proximity; and (2) Proper droplet formation, which can be hampered by extremely long DNA molecules. Digestion was performed in 10 μl including 2.5 U of restriction enzyme, 1x of the corresponding buffer, and 250 ng of genomic DNA at 37 °C for 3 hours, and then 2 μl (50 ng) were directly used as template in the ddPCR reaction. Since DNA molecule size reduces over time, ddPCR genotyping was performed by decreasing order of IR size, starting with the inversions with higher DNA quality requirements.

There were several reasons why a ddPCR result was not valid: (1) Total linkage >7.5% was required to be considered reliable, which was only an issue in those inversions with the largest IRs; (2) Droplet count below 10,000 due to the presence of undigested high-molecular-weight DNA; (3) Intermediate linkage ratios between the expected values for homozygotes and heterozygotes (e.g. ∼0.25 or ∼0.75); and (4) Small deletions or duplications of one or more of the amplicons that result in ratios between them that were different than 1 in certain individuals (Supplemental Table S3). Except for the altered amplicon ratios, in most cases these problems were solved by repeating the ddPCR reactions using a fresh DNA dilution from stock or storing the diluted DNA at 4 °C for a few days.

### Genotype clustering

Inversion genotypes were called by clustering all *O1* linkage ratios (Supplemental Fig. S5), except for those based on a total linkage <7.5%, or <15% if only one measurement was available. Also, in HsInv0382, where a deletion increases the linkage in one of the breakpoints, samples genotyped only by one breakpoint were excluded. For each inversion, we calculated the euclidian distance between individuals (stats::dist R function) (R Core Team 2017) using two randomly-selected *O1* linkage ratios per sample scaled to normal scores (base::scale R function). Since in some cases there is a variable number of measurements, this process was repeated 200,000 times and a mean pairwise distance between individuals was obtained. We performed a hierarchical clustering analysis (ward.D implemented method) on this similarity matrix to determine group membership (stats::hclust, stats::cutree R functions) (R Core Team 2017). Clustering was run to find two or three clusters that were defined taking into account that heterozygotes should be centered around 0.5. To assess the uncertainty of sample classification, we clustered two thirds of our samples selected at random 10,000 times and their percentage of association to each genotype was measured. Individuals that appeared as outliers in the membership distribution and were included >5% of times in a different cluster were not genotyped. For chr. X inversions, we repeated the analysis only with males clustered into *O1* or *O2* and, if these genotypes were more robust, they were the ones used. Finally, we tried to recover samples without a clear genotype in an extra clustering step by selecting proportionally more often those *O1* linkage ratios based on a higher total linkage when calculating euclidean distances and repeating the bootstrapping to obtain the final genotype clusters.

### Haplotype fusion PCR (HF-PCR)

HF-PCR was carried out in an oil and water emulsion to generate the fused amplification product followed by a regular PCR with nested primers (Supplemental Table S15). To separate both inversion breakpoints, genomic DNA was digested overnight at 37 °C in a volume of 20 μl including 5 U of SwaI for HsInv395 or SalI for HsInv605, 1x buffer and 250 ng DNA, the restriction enzyme was then heat inactivated at 65 °C for 15 min, and 25 ng of DNA were used as template. Emulsion PCR reactions were performed in 25 μl in 96-well plates for 40 cycles as previously described (Turner and Hurles 2009). The main differences were that we used SOLiD^TM^ EZ^TM^ Bead Emulsifier Oil Kit (Applied Biosystems) to form the emulsions, and that after amplification we carefully transferred the emulsion to a fresh plate, added 50 μl of 1x Phusion HF buffer (Thermo Scientific) to increase volume, centrifuged for 5 minutes at maximum speed and recovered the aqueous phase containing the amplification products. Next, we did a 30-cycle reamplification step with 1 μl of a 1/10 dilution of the previous PCR, 1.5 U Taq DNA polymerase (Roche), and 200-400 nM of each of the three nested primers in a 25 μl total volume (Turner et al. 2006). Finally, 10 μl of the PCR reaction were loaded into a 3% agarose gel for visualization.

### Analysis of inversion frequency

Frequency differences between populations were measured with Weir and Cockerham’s F_ST_ estimator implemented in vcftools (v0.1.15) (Danecek et al. 2011), using the 92 samples common to the 1000GP and only females for chr. X inversions or paired male chromosomes for the chr. Y inversion. F_ST_ values of each inversion were compared with an empirical distribution from 10,000 genome-wide biallelic 1000GP SNPs polymorphic in at least two of the populations, matched by chromosome type (autosome or chr. X) and excluding those SNPs overlapping inversion regions ±100 kb. Correlation between MAF and the logarithm of the physical and genetic lengths of inversions was measured with a linear model implemented in robustbase::lmrob R function (Maechler et al. 2018), including data from 45 inversions in a larger sample of the same populations (Giner-Delgado et al. 2019). Inversion physical length corresponds to the distance between IRs and genetic length was interpolated from Bhérer at al. (2017) (Bhérer et al. 2017) recombination map, using the female map for chr. X and the sex average map for autosomes. No genetic length was available for chr. Y inversions and HsInv0608 (chr. X), which falls outside the last marker in the map.

### Linkage disequilibrium (LD) analysis

Pairwise LD between genotypes of inversions and neighboring biallelic single nucleotide changes and small indels from 1000GP Phase 3 (inversion region ±1 Mb) was calculated using the *r^2^* statistic with PLINK v1.9 (Purcell et al. 2007) separately for each population group and the 92 samples common to both datasets. SNPs were further classified as shared, private or fixed, depending on whether they were unambiguously polymorphic in the two orientations, polymorphic only in one, or their alleles were in perfect LD with the inversion, respectively (Giner-Delgado et al. 2019). To minimize possible genotyping errors in 1000GP data, we based our analyses on reliable variants, defined as those located in accessible regions according to the 1000GP strict accessibility mask and that do not overlap known SDs (Bailey et al. 2002).

### Recurrence analysis

To generate haplotypes of the inverted regions, we selected the same 1000GP reliable variants used in the previous analysis that were present in at least two genotyped chromosomes. When the number of variants in an inversion was below 50, those accessible according to the pilot criteria were also included to maximize information. Available 1000GP Phase 3 haplotypes were used as scaffolds and the phase of the inversion genotypes was inferred using MVNcall (Menelaou and Marchini 2013) by positioning inversions in the middle as a single-nucleotide variant. Haplotypes were clustered by similarity and their relationships were visualized using the iHPlots strategy (Giner-Delgado et al. 2019). The putative ancestral orientation and the original inversion event were defined based on the *O1* and *O2* haplotype diversity and the frequency and geographical distribution of haplotypes (usually considering as ancestral those found in African samples) (Supplemental Table S8). Additional independent inversion or re-inversion events were identified conservatively as haplotype clusters with an unexpected orientation and clearly differentiated from other *O1* or *O2* haplotypes (≥3 differences spanning >3 kb that cannot be easily explained by a recombination event) (Fig. 4B-C). To avoid possible phasing errors, in inversion heterozygotes we tested whether switching the orientation of both haplotypes still supported recurrence or not. For HsInv0416 we used the known chr. Y haplogrup information to determine the number of inversion events (Poznik et al. 2016) and estimated the recurrence rate as in Giner-Delgado et al. (2019) (Giner-Delgado et al. 2019). Briefly, a number of 30,931.1 generations were calculated for all branches involved in the phylogeny that relates the 48 analyzed males (Poznik et al. 2016), including a C-T branch split time of 76,000 years, a total number of mutations of 5,591, and an average number of mutations of all branches of 549.5, plus a generation time of 25 years (Repping et al. 2006). Finally, we tested different variables to determine their effect on the number of recurrent events per chromosome: chromosome type (autosome, chr. X or chr. Y), inversion and IR length, IR/Inv size ratio, IR identity, and PRDM9 motifs/kb within IRs (Myers et al. 2008). The model was built by stepwise regression with forward selection using the robustbase::lmrob R function (Maechler et al. 2018) and logarithm-transformed values to remove outliers.

### Gene expression analysis

Inversion effects on LCL gene expression were first analyzed in 30 CEU and 29 YRI experimentally-genotyped individuals from the Geuvadis project (Lappalainen et al. 2013). We excluded HsInv0416 in chr. Y due to the low statistical power to detect differences only in males. Inversions 17p21, HsInv0290, HsInv0395, HsInv0605 and HsInv0786 were also imputed in 328 additional Geuvadis European samples (59 CEU, 91 TSI, 86 GBR and 92 FIN) using a representative tag SNP in the CEU population (which except for HsInv0290 belongs to a larger set of SNPs in LD in the expanded population). HsInv1057, with tag SNPs in YRI, was imputed in 58 additional Geuvadis YRI individuals but no significant gene-expression associations were obtained. RNA-seq reads (EMBL-EBI ArrayExpress experiment E-GEUV-1) were aligned against the human reference genome GRCh38.p10 (excluding patches and alternative haplotypes) with STAR v2.4.2a (Dobin et al. 2013). We estimated gene expression levels as reads per kilobase per million mapped reads (RPKM) based on GENCODE version 26 annotations (Harrow et al. 2012) and quantified transcript expression with RSEM v1.2.31 (Li and Dewey 2011), filtering out non-expressed genes and transcripts with <0.1 RPKM in >80% of the samples. RPKM values were normalized by quantile transformation across all samples and expression of each gene/transcript was adjusted to a standard normal distribution by rank-based inverse normal transformation. Association with expression of 418 genes and 2,044 transcripts was calculated for all biallelic variants with MAF >0.5 (including the inversion) within 1 Mb at either side of the transcription start site using linear regressions implemented in FastQTL (Ongen et al. 2016). Since technical or biological confounders reduce the power to find associations, we adjusted expression values by the top three 1000GP genotyping principal components (corresponding to population structure), sequencing laboratory, gender, and an optimal number of PEER (probabilistic estimation of expression residuals) components (Stegle et al. 2012) for eQTL finding (for genes and transcripts, respectively, 12 and 15 for the experimental and 25 and 30 for the imputed set). Significant INV-eQTLs correspond to a *Q* value false-discovery rate (FDR) <0.05 (Storey & Tibshirani 2003).

Next, we estimated inversion gene-expression effects in other tissues through the LD between inversion alleles and eQTLs in GTEx V7 release (GTEx Consortium 2013, 2017) using FAPI v0.1 (Kwan et al. 2016). First, we randomly took three samples of 30 experimentally genotyped individuals following ethnic proportions of GTEx donors (25 individuals from CEU, 4 YRI and 1 EAS) per inversion-gene pair and tissue to calculate LD patterns between each inversion and neighbouring SNPs, which were subsequently used to impute the corresponding inversion association *P* values from GTEx eQTL *P* values. If any *P* value was lower than the genome-wide empirical threshold defined by GTEx for each gene and tissue (GTEx Consortium 2017), we generated 30 samples of 30 individuals to calculate statistical significance more accurately. The eQTL *P* value of the inversion was defined as the median of permuted *P* values and the association confidence interval as the 25th and 75th percentiles, since the small number of individuals for LD calculation can produce extreme *P* values. In addition, we filtered out those associations with estimated *P* value lower than GTEx significance threshold or with a confidence interval spanning more than two orders of magnitude. The conservative nature of this analysis is represented by HsInv0389, in which only four of the 16 genes previously associated to the inversion based directly on the LD with GTEx eQTLs from a larger number of samples were identified. For both Geuvadis and GTEx, the most significant associated variants for each gene/transcript were designated as lead eQTLs. Moreover, inversions in high LD with the top marker (*r^2^* ≥ 0.8) were indicated as potential lead eQTLs. GTEx LD was estimated as the median LD of permutations as explained above. Effect sizes were calculated as a function of MAF and *P* value, whereas direction was determined through LD with eQTLs using PLINK v1.9 *--r2 in-phase* option (Purcell et al. 2007).

### GWAS data analysis

The impact of inversions in relevant phenotypic traits was assessed with the GWAS Catalog curated collection of the most significant SNPs associated to a particular phenotype (*P* < 10^-5^) (http://www.ebi.ac.uk/gwas/) [release 2018-06-25, v1.0] (MacArthur et al. 2017). First, to explore the enrichment of GWAS SNPs within inversions, we translated GWAS Catalog coordinates to hg19 using Ensembl REST API (Yates et al. 2016) and grouped together the signals associated to SNPs in high LD (*r^2^* ≥ 0.8) in 1000GP data and corresponding exactly to the same phenotype, resulting in 67,035 non-redundant SNP-trait associations. Then, we created a background distribution of each inversion with 1,500 random genomic regions of the same size that the inverted segment to calculate the enrichment *P* values. We excluded from permutations chr. Y, gaps, and the major histocompatibility complex region (chr6:28,477,797- 33,448,354), known to harbor a vast number of associations. In addition, we tested that the GWAS enrichment was not biased by the allele SNP frequencies by selecting 150 random regions per inversion with comparable patterns of common variants (number of 1000GP loci with global MAF > 0.05 per kb ±20%) and without this criteria, which showed very similar results (*r^2^*= 0.99). To explore which inversions were driving the enrichment, we repeated the analysis for each inversion independently using a one-tailed permutation test (to account for inversions with zero GWAS signals). Also, we compared the proportion of genes related to particular clinically relevant traits or diseases as reported in the GWAS Catalog inside or around (±150 kb) each inversion with respect to the rest of the genome (Supplemental Table S11). Only traits with at least four associated genes close to the inversion were considered and *P* values were calculated with a Fisher’s exact test adjusted by Bonferroni correction. Finally, we crossed GWAS Catalog variants with those in high LD with our inversions (*r^2^* ≥ 0.8) in the corresponding population or the closest one available in our data, while the global LD was used for GWAS with populations from different continents (Supplemental Table S12).

## DATA ACCESS

All inversion genotypes are available at InvFEST (http://invfestdb.uab.cat) and have been deposited in NCBI’s dbVar database of genomic structural variation (https://www.ncbi.nlm.nih.gov/dbvar) under accession number XXXXX.

## ACKNOWLEDGEMENTS

We would like to thank Salvador Bartolomé, Xavier Alba and James Thomas for technical support and advice, Alba Vilella, Marina Laplana and Sergi Villatoro for help with DNA isolations and microsatellite analysis, and Xavier Estivill and Marta Morell for the European and African lymphoblastoid cell lines. This work was supported by research grants BFU2013-42649-P and BFU2016-77244-R funded by the Agencia Estatal de Investigación (AEI, Spain) and the European Regional Development Fund (FEDER, EU), ERC Starting Grant 243212 (INVFEST) from the European Research Council under the European Union Seventh Research Framework Programme (FP7), and 2017-SGR-1379 from the Generalitat de Catalunya (Spain) to MC, and a La Caixa Doctoral fellowship to JLJ. MGV was supported by POCI-01-0145-FEDER-006821 funded through the Operational Programme for Competitiveness Factors (COMPETE, EU) and UID/BIA/50027/2013 from the Foundation for Science and Technology (FCT, Portugal). ddPCR reagents used in this study were provided by Bio-Rad Laboratories, Inc.

## AUTHOR CONTRIBUTIONS

MP and MC designed the genotyping assays and oversaw all steps; MP, SP, DI and AD performed experiments; CGD, MGV, MP and MC analyzed evolutionary data; JLJ and MP analyzed functional effects; MP, SP, JFR, GKN and MC contributed to ddPCR assay design and optimization; MP, JLJ and MC wrote the paper.

